# Downregulation of Transducin Delays Photoreceptor Degeneration in P23H Rhodopsin Retinitis Pigmentosa

**DOI:** 10.1101/2025.10.04.680427

**Authors:** Deepa Mathew, Lee Sturgis, Marianna Mathison, Frans Vinberg

## Abstract

In inherited blinding diseases, such as Retinitis Pigmentosa (RP), photoreceptors progressively degenerate, eventually leading to blindness. Unfortunately, effective treatments to prevent or delay vision loss do not exist for most RPs. Dark rearing is known to delay retinal degeneration in preclinical models of RP. Therefore, in this study we evaluated the impact of reducing photoreceptor light signaling on RP progression. This was done by genetically ablating or downregulating transducin in rods or cones in a preclinical RP model carrying a single P23H mutant rhodopsin allele (P23H mice). Ablating rod transducin significantly improved photoreceptor survival in the P23H retina. Additionally, downregulating rod transducin promoted photoreceptor survival and improved rod light response in P23H mice. Remarkably, male P23H mice retained robust cone function until old age in the absence of rod transducin whereas female P23H carriers experienced significantly faster loss of cone function. In these females, reducing cone transducin improved cone function whereas the same treatment was not effective in male P23H carriers. Our data demonstrate that reducing rod or cone transducin expression in P23H mice improves the survival and function of rods and cones, and suggest transducin downregulation as an effective therapeutic strategy to delay photoreceptor degeneration in RP.

## INTRODUCTION

Progressive visual impairment and blindness is a major health concern necessitating the development of new therapeutic approaches. One of the main causes of vision loss is inherited retinal degenerations such as retinitis pigmentosa (RP), which mostly involves progressive loss of rod photoreceptors followed by secondary loss of cones (1). RP is caused by mutations in genes affecting the normal structure, function, or metabolism of rod photoreceptors or the retinal pigment epithelium (RPE)(1, 2). Till date there are no cures for RP, other than a few gene therapy approaches showing promising results (3). However, the heterogeneous disease etiology of RP makes targeted gene therapy impractical and expensive. Therefore, it is important to develop gene agnostic therapies for delaying rod degeneration and preserving cone function so that patients can maintain useful vision during RP progression.

A subset of RP is caused by mutations of rhodopsin, the light-sensitive protein in the rod photoreceptor cells of the retina which is crucial for dim light vision. Among them the P23H mutation of rhodopsin is the most common cause of autosomal dominant retinitis pigmentosa (adRP) in the USA. This mutation has been studied extensively in a variety of experimental systems providing insights into its pathogenic mechanisms (3). The P23H mutation affects overall structure and activity of rhodopsin including chromophore binding, stability and activity of the monomer and its dimerization preference (4, 5). The mutant rhodopsin causes unfolded protein responses leading to Endoplasmic Reticulum (ER) stress and mitochondrial dysfunction and exerts a destructive effect on disc membrane morphogenesis (3, 6–8). Better understanding of these different pathogenic mechanisms and cellular response pathways against them during RP progression will aid in developing novel gene agnostic therapies.

In some preclinical models of RP, disease progression is accelerated by bright light and/ or slowed by dark rearing (9). Light activation of rhodopsin contributed to the severity of the degeneration in animal models of P23H rhodopsin and dark rearing was protective in some of these studies (10–12). During rod phototransduction, light is converted into electrical signals by a series of steps initiated by the isomerization of 11-cis retinal bound to rhodopsin. The resulting activated rhodopsin further activates its G protein Transducin α which effectively propagates the light signaling leading to the hyperpolarization of rod photoreceptor membrane (13). Here, we investigated the effect of inhibiting light signaling by downregulating transducin expression as a therapeutic strategy to delay RP progression, using a mouse model of P23H rhodopsin mutation (P23H mice) that closely recapitulates slow progressing adRP in patients (6). For that we genetically manipulated the function of rod transducin (*Gnat1*) or cone transducin (*Gnat2*) in P23H mice and evaluated the effectiveness of this strategy by assessing retinal structure and function during disease progression. Since females display accelerated photoreceptor degeneration in these mice, we carefully compared the impact of manipulating transducin expression on RP progression between sexes (14, 15).

## RESULTS

### Elimination of rod transducin delays photoreceptor degeneration in P23H mouse model of autosomal dominant retinitis pigmentosa

Previous studies have reported that increased light exposure exacerbates retinal degeneration while dark rearing delays the progression of degeneration in animal models of RP (10, 11). Similar to dark rearing, inhibiting rod phototransduction by abolishing Gnat1 has protective effect in some forms of RP but not in all RP models (16, 17). The effect of modifying rod phototransduction in slowly progressing RP caused by rhodopsin mutations has not been studied before. To address this gap in knowledge, we used both sexes of the well-established adRP mouse model, P23H mice (*Rho^WT/P23H^*), and genetically ablated *Gnat1* in them (18) to block the rod light response. We used Optical Coherence Tomography (OCT) imaging to assess retinal structure and quantified Outer Nuclear Layer (ONL) thickness which represents the number of surviving photoreceptor nuclei. At 7 months, the ONL thickness was reduced to 23% in P23H males and 11% in P23H females compared to their WT (C57Bl/6J) littermates (Figure1A, 1B). This accelerated degeneration in females was consistent with the previous reports (14, 15). Remarkably, rod survival was dramatically improved in the absence of rod transducin as demonstrated by increase in ONL thickness by 86% and 196% in *P23H Gnat1^-/-^* males and females respectively, as compared to *P23H Gnat1^+/+^* littermates, (Figure 1A, 1B). Interestingly, WT mice showed a 10 % reduction in ONL thickness when *Gnat1* was abolished, suggesting a slow degeneration in these mice, (Figure 1B). At 12 months, P23H mice with WT transducin levels showed the characteristic fundus appearance of advanced retinitis pigmentosa (Figure S1). This was absent even at 12 months when Gnat1 was completely abolished reflecting the protective effect of eliminating light signaling in rods during RP progression.

**Figure 1.**
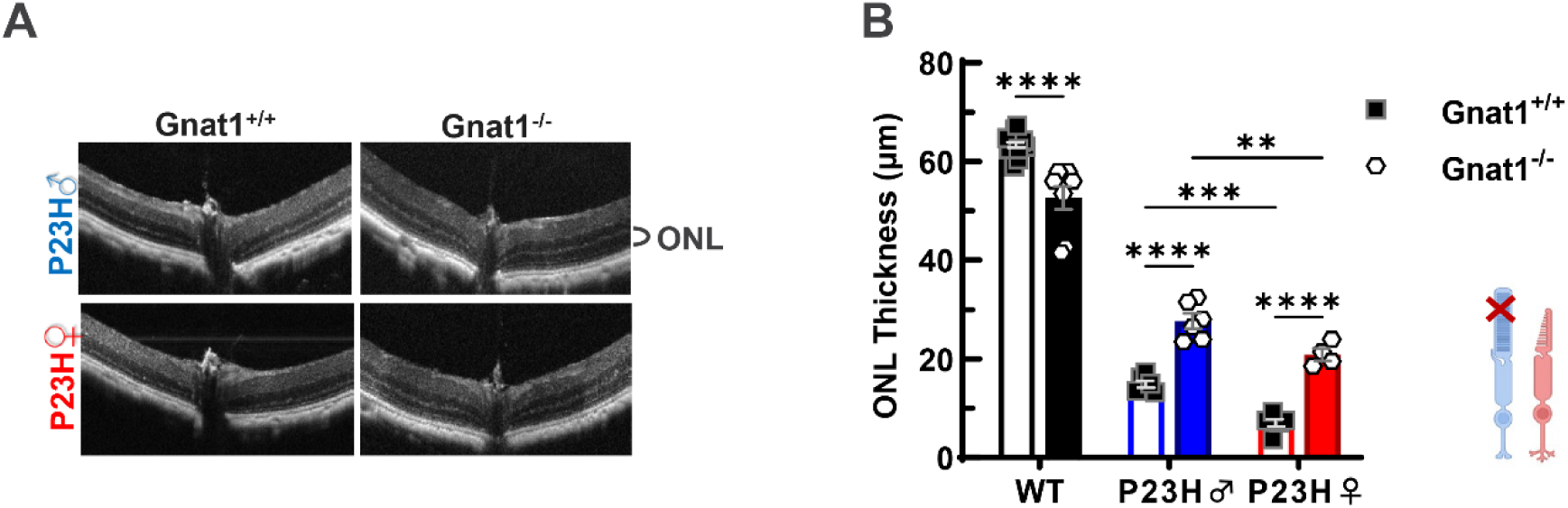
Abolishing rod transducin delays photoreceptor degeneration in P23H mice. Representative OCT images of retinas from 7-month-old *P23H Gnat1^+/+^* and *P23H Gnatl^7^-* males and females (A). Quantification of ONL thickness from the central retina from WT, P23H male and P23H female mice with either normal *Gnatl* expression or *Gnatl* ablated (B) showing improved photoreceptor survival in the P23H retinas when *Gnatl* is ablated (N=5-13). Tukey’s post hoc tests: **p<0.01, ***p<0.001, ****p<0.0001.

### Reducing rod transducin delays photoreceptor degeneration and improves rod function in P23H model of adRP

Abolishing rod phototransduction significantly delayed photoreceptor degeneration in P23H mice. However, it would not be therapeutically easy or advisable to completely abolish *Gnat1* expression as it would be expected to lead to night blindness. Downregulation of transducin, on the other hand, would be feasible and is not expected to severely compromise rod function. It is not known, however, if partial suppression of rod phototransduction by reducing rod transducin expression has any effect on RP progression. Rod transducin expression was reduced by 25% in heterozygous Gnat1 mice (*Gnat1^+/-^*) (19). Therefore, to evaluate the effect of downregulating phototransduction on photoreceptor degeneration we used *P23H Gnat1^+/-^* mice and assessed their retinal structure using OCT imaging. These mice were bred in the absence of cone transducin (*Gnat2^-/-^*) to eliminate cone light response and isolate rod function in the subsequent experiments. Ablating Gnat2 did not show any effect on photoreceptor survival in P23H mice (Figure S2). At 3 months, the ONL thickness was reduced to 42% in *P23H Gnat2^-/-^*males and 35% in *P23H Gnat2^-/-^* females compared to *WT Gnat2^-/-^* (Figure 2A, 2B) consistent with the previous reports of faster RP progression in females (14, 15). In these mice reducing rod transducin levels resulted in 43% and 34% improvement of ONL thickness in P23H males and females, respectively, compared to their counterparts with normal *Gnat1* expression (Figure 2A, 2B). On the contrary, WT mice showed a small but significant 7.5% reduction in the ONL thickness when *Gnat1* was heterozygous (Figure 2B). The markedly improved photoreceptor survival in P23H mice by partial suppression of *Gnat1* expression points to the possibility of targeting *Gnat1* as a strategy to delay degeneration in RP.

**Figure 2.**
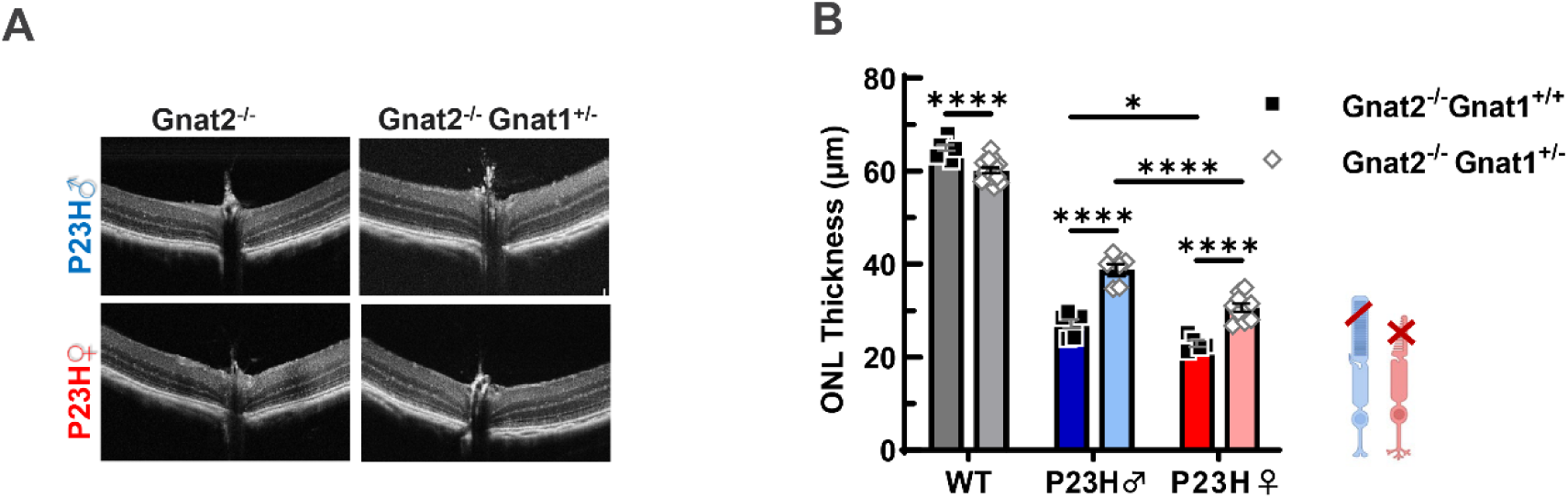
Reducing rod transducin delays degeneration in P23H Retina. Representative OCT images of retinas from 3-month-old *P23H Gnat2^ Gnat1^+/+^* and *P23H Gnat2^/^-Gnat1^+/^∼* males and females (A). Quantification of ONL thickness analysis from the central retina of WT, P23H male and P23H female mice (B) showing improved photoreceptor survival in the P23H retinas in *Gnatl* heterozygous conditions. N=5-13. Tukey’s post hoc tests: *p<0.05, **p<0.01, ***p<0.001.

As a possible therapeutic strategy, it is important to validate functional preservation of photoreceptors with reduced rod transducin in P23H mice. To evaluate retinal function, we performed transretinal electroretinogram (ex vivo ERG) on isolated WT and P23H retinas with either *Gnat1^+/+^* or *Gnat1^+/-^* condition and pharmacologically isolated photoreceptor response and bipolar cell response to light flashes of increasing intensity. These mice also lacked cone transducin (*Gnat2^-/-^*) abolishing cone phototransduction and ensuring the isolation of rod response. As expected, rod light response was reduced in 3 month old P23H *Gnat2^-/-^*males and females compared to WT (Figure S3). We did not observe any significant difference in rod function between P23H males and females at 3 months on the contrary to the previous reports of faster loss of total ERG response in females and irrespective of the reduced ONL thickness in females (Figure 2B). When rod transducin was reduced, P23H *Gnat2^-/-^ Gnat1^+/-^* males and females showed significant improvement in rod response amplitude (Figure 3A, 3C). Such improvement in function was also maintained in rod mediated bipolar cell response amplitude when *Gnat1* expression was reduced (Figure 3B and 3D). As expected, when the rod transducin expression was normal, the maximum rod response amplitude (Rmax) was reduced to 40% in both P23H *Gnat2^-/-^*males and females compared to WT *Gnat2^-/-^*. However, when transducin expression reduced, Rmax improved by 49% and 55% in *P23H Gnat2^-/-^ Gnat1^-/-^* males and females respectively compared to those with normal rod transducin expression (Figure 3E). The light sensitivity of rods, determined based on half-saturating intensity (I_1/2_), was reduced in *P23H Gnat2^-/-^*males and females compared to WT *Gnat2^-/-^* consistent with previous reports (20). When rod transducin expression was reduced in *P23H Gnat2^-/-^* mice, the mean rod sensitivity was 19% and 24% improved in males and females respectively, though this improvement was not statistically significant (Figure 3F).

**Figure 3.**
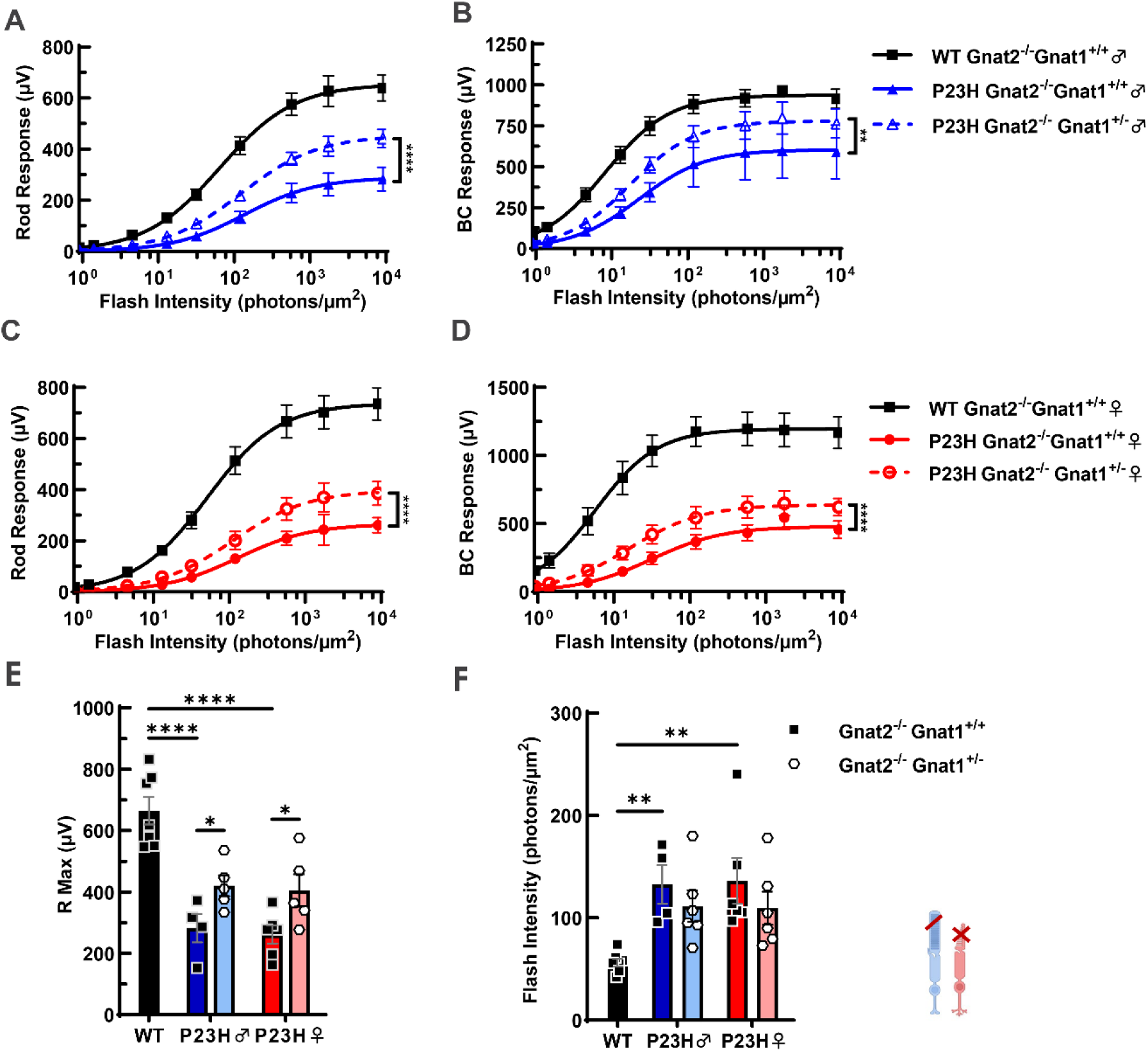
Reducing rod transducin improves rod function in P23H Retina. Rod light response amplitude as a function of light flash intensity from 3-month-old *WT Gnat2’* and *P23H Gnat2’* males and females with either *Gnat1^+/+^* or *Gnat1^+/^* conditions. Rod response (A) and rod mediated bipolar response (B), from males; rod response (C) and rod mediated bipolar cell response (D) from females demonstrating improved rod function when *Gnatl* expression was reduced. Mean + SEM; two-way ANOVA, **p<0.01, ****p<0.0001. Smooth traces in (A-D) are Hill’s equation fitted to the mean amplitude data. Comparison of maximum rod response amplitude (E) and intensity required to generate half-maximal rod response (F). Mean ± SEM, N=4-7, Tukey’s post hoc tests, *p<0.05, **p<0.01, ****p<0.0001.

In the WT mice reducing Gnat1 expression did not alter rod light response (Figure S4A) even though these mice had reduced ONL thickness (Figure 2B). Interestingly, reducing *Gnat1* expression resulted in significant reduction in rod mediated bipolar cell response in WT *Gnat2^-/-^* (Figure S4 B) suggesting changes in rod to rod bipolar synaptic transmission when rod transducin is reduced. Since our ERG recording was performed in dark adapted condition, it rules out the possibility of transducin translocated to rod synaptic spherules in response to light, modulating rod to rod bipolar cell synaptic transmission as reported previously (21). However, the possibility of reduced transducin expression or translocation leading to stable changes at the rod synapse cannot be overlooked.

#### Cone function is well-preserved in P23H male mice when rod transducin is abolished

In most RP cases disease progression starts with rod photoreceptor degeneration and associated loss of night vision. Prolonging cone function in such cases will help preserve essential color and high acuity vision in patients. To understand the mechanisms of cone cell death in RP for identifying better therapeutic strategies, we investigated progression of cone function loss in the degenerating P23H retina using ex vivo ERG. Rod transducin/rod light signaling was ablated (*Gnat1^-/-^*) in these mice ensuring the isolation of cone responses. Surprisingly, we found that the cone light responses were better preserved in males while females showed more severe loss of cone function at this age (Figure 4A). This preservation of function was also observed in the cone mediated bipolar cell response (Figure 4B). Better survival of photoreceptors in P23H mice when *Gnat1^-/-^* is ablated, might also be contributing to the preserved cone function in these mice (Figure 1A, 1B). The maximum cone response amplitude was maintained at 67% in *P23H Gnat1^-/-^* males while it was reduced to26% in *P23H Gnat1^-/-^* females compared to *WT Gnat1^-/-^* mice (Figure 4C). The cone light sensitivity (I_1/2_) was increased by 10% in *P23H Gnat1^-/-^* males and reduced by 22% in *P23H Gnat1^-/-^* females compared to WT *Gnat1^-/-^*, though these changes were not statistically significant (Figure 4D). Though previous studies have reported sex hormone dependent accelerated photoreceptor degeneration in P23H females (14, 15), our data demonstrated the sex difference specifically in cone function.

**Figure 4.**
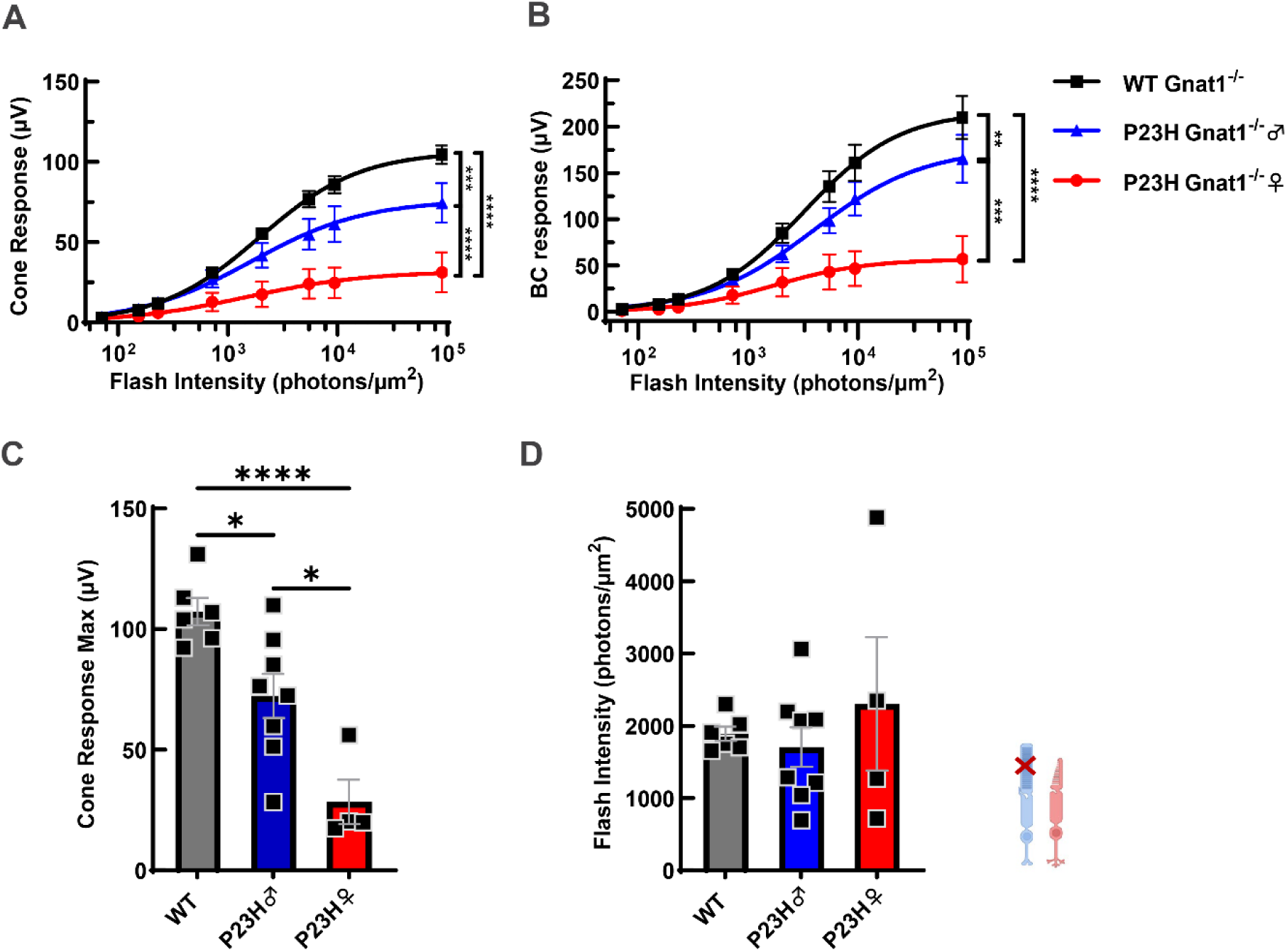
Cone function in male *P23H Gnatl∼^1^*is well preserved at 12-14 months. Ex vivo ERG amplitude data as a function of light flash intensity from 12-14-month *WT GnatT^7^’* and *P23H GnatT^7^’* males and females showing cone response (A), and cone mediated bipolar cell response (B), demonstrating better preserved cone function in males and accelerated loss of function in females. Smooth traces in (A and B) are Hill’s equation fitted to the mean amplitude data. Two-way ANOVA, **p<0.01 ***p<0.001, ****p<0.0001. Comparison of maximum cone response amplitude (C) and cone light sensitivity determined as the intensity required to generate half-maximal cone response (D). Tukey’s post hoc tests, *p<0.05, ****p<0.0001. Mean ± SEM, n=4-8

### Reducing cone transducin expression rescued cone function in P23H female mice

Female cone function declined significantly by 1-year of age in the P23H mice. Since reducing rod transducin expression was protective to rod function, we investigated if reducing cone transducin (*Gnat2*) expression has any effect on cone degeneration in P23H retina. We recorded cone responses using ex vivo ERG from littermates of *P23H Gnat1^-/-^* mice from Figure 4, with one *Gnat2* allele deleted (*P23H Gnat1^-/-^ Gnat2^+/-^*) at 12-14 months. In these mice, with reduced cone transducin expression, female cone function was markedly improved compared to female *P23H Gnat1^-/-^* with normal *Gnat2* expression (Figure 5A). Consistently, cone mediated bipolar cell response was also significantly improved in these females (Figure 5B). The maximum cone response amplitude was increased by 1.9 fold in *P23H Gnat1^-/-^ Gnat2^+/-^* females compared to those with normal *Gnat2* expression, though this increase was not statistically significant (Figure 5C). unlike in females, downregulation of *Gnat2* was not protective in P23H males (Figure S5 A-D). On the contrary, in WT mice, downregulation of *Gnat2* led to a significant reduction of cone light response and cone mediated bipolar cell response at 12-14 months of age (Figure S6A, S6B). The maximum cone response amplitude and cone light sensitivity were also reduced in *WT Gnat1^-/-^ Gnat2^+/-^* mice (Figure S6C, S6D). Thus, reducing *Gnat2* in cones had different effects in WT, P23H males and females showing a genotype and sex dependent effect of downregulating *Gnat2* on cone function.

**Figure 5.**
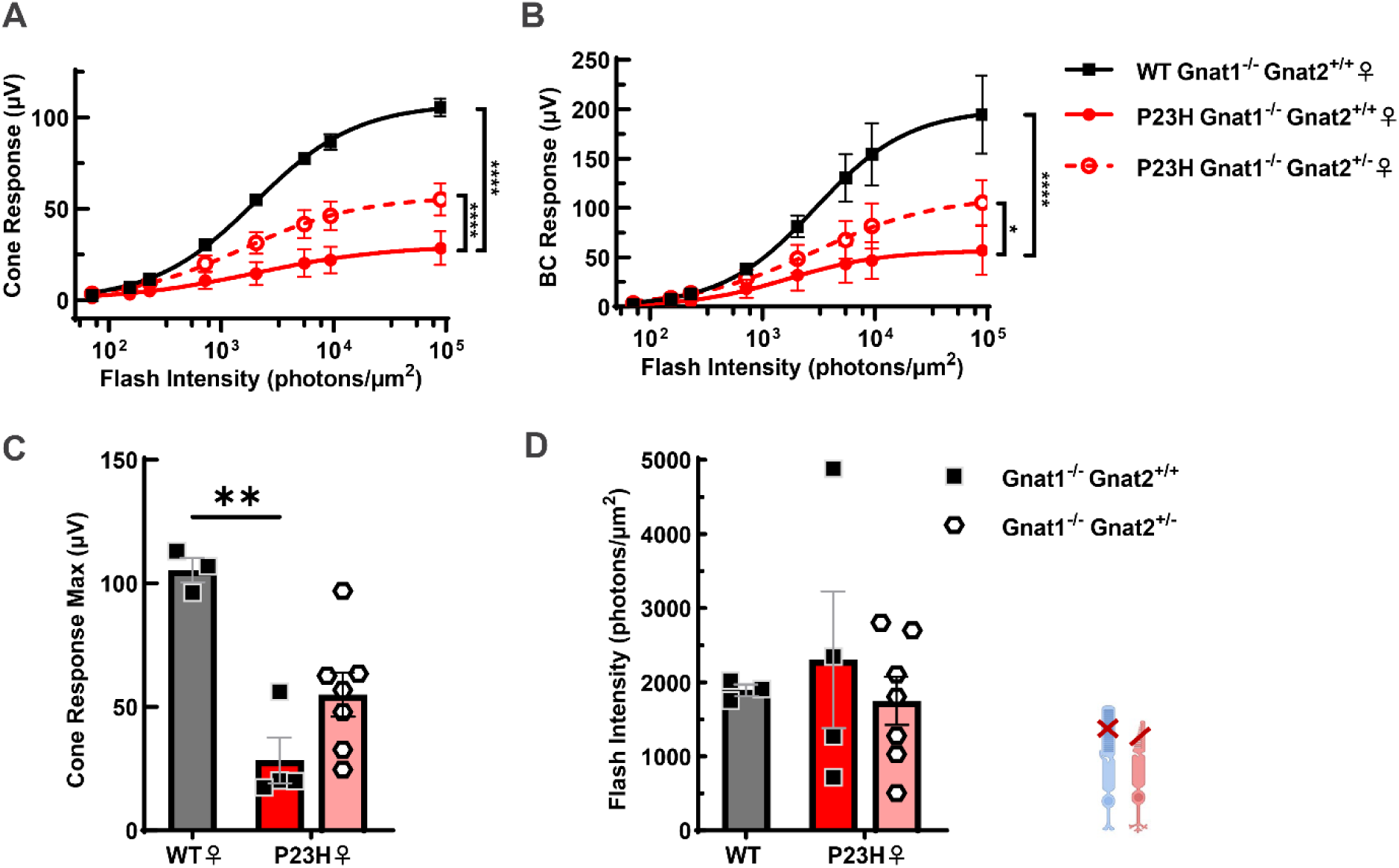
Reducing cone transducin is protective in P23H females. Cone light response to light flashes of increasing intensities recorded using ex vivo ERG from 12-14 months old *WTGnatl’^7^’, P23H GnafH-Gnat2^+/+^* and *P23H Gnatl’^1^’-* Gnaŕ2^+/^-females showing cone photoreceptor response (A) and cone mediated bipolar cell response (B) demonstrating the protective effect of reducing *Gnat2* expression. Smooth traces in (A and B) are Hill’s equation fitted to the mean amplitude data. Two-way ANOVA, *p<0.05, ****p<O.0001. Comparison of maximum cone response amplitude (C) and intensity required to generate half-maximal cone response (D). Tukey’s post hoc tests, **p<0.01. Mean ± SEM, N=3-7

### Reducing cone transducin did not affect photoreceptor survival in P23H mice

Since we observed sex and genotype specific changes in cone function when Gnat2 was heterozygous, we evaluated photoreceptor survival by quantifying ONL thickness from OCT images. In *WT Gnat1^-/-^ Gnat2^+/-^* mice with reduced cone function (Figure S6), we did not observe the ONL thickness to be significantly different from those with normal *Gnat2* expression, suggesting that the total photoreceptor number is comparable between these groups (Figure 6A). Similarly reducing cone transducin in *P23H Gnat1^-/-^* males and females did not improve photoreceptor survival (Figure 6A). Additionally, we quantified the number of cone photoreceptors from the confocal images of peanut agglutinin (PNA) stained sections of the retina (Figure S7). Cone photoreceptor survival was not significantly altered when *Gnat2* was reduced in *WT Gnat1^-/-^*or *P23H Gnat1^-/-^* mice (Figure 6B). These observations do not explain the changes in cone function observed in these mice. However, we did not quantify any possible changes in cone outer segment length in *Gnat2^+/-^* condition. In some *WT Gnat1^-/-^ Gnat2^+/-^* mice OCT imaging showed retinal detachment (data not shown) which may also contribute to the loss of cone function. Overall, the improvement of cone function in P23H females by reduced *Gnat2* expression does not seem to be via improved rod or cone survival but something intrinsic to the cone photoreceptors.

**Figure 6.**
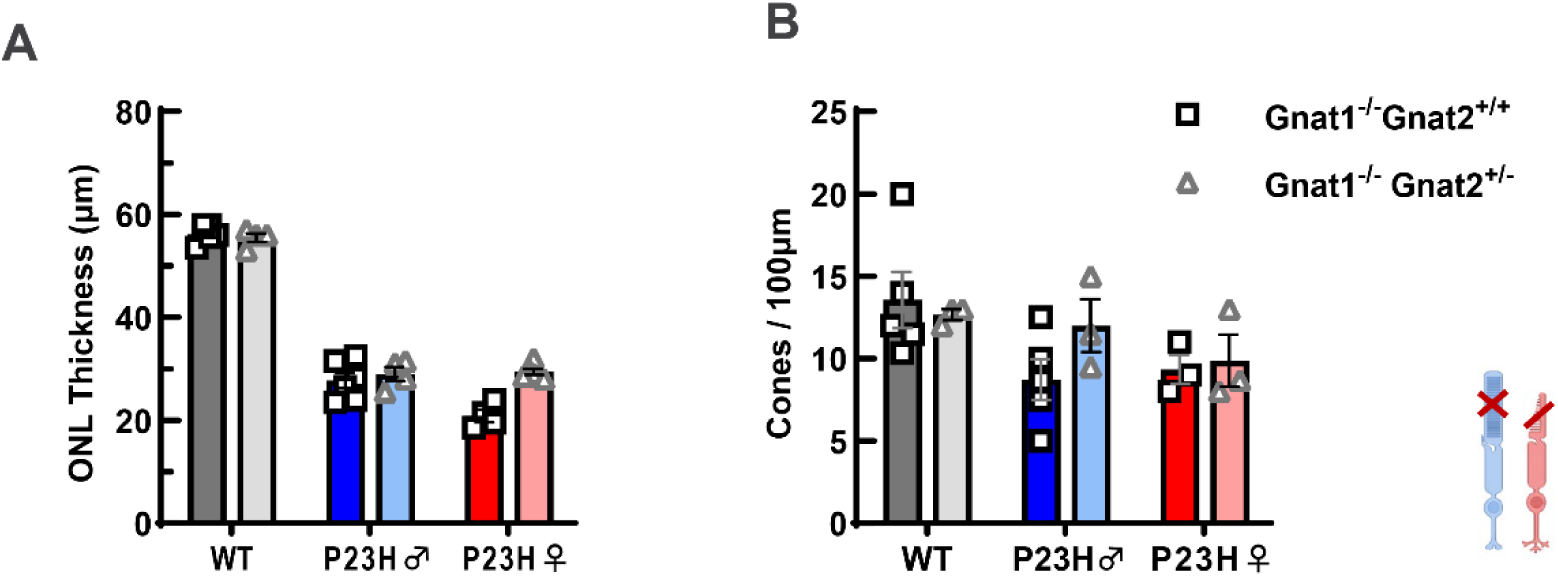
Reducing *Gnat2* did not affect photoreceptor survival in P23H retina: Quantification of ONL thickness from 7 months old *WT Gnatl’^7^’* and *P23H Gnatl^^7^’* males and females compared with those harboring a single *Gnat2* allele (A) showing no significant effect of Gnat2 expression on photoreceptor number. Quantification of cone photoreceptors stained with peanut agglutinin from retina sections of *P23H Gnatl’^7^’* and *P23H Gnatl’^7^’with* normal or reduced *Gnat2* expression (B) showing no significant effect of *Gnat2* heterozygous condition on cone survival. N=3-6

## DISCUSSION

### Reducing transducin; a gene agnostic therapeutic approach

Therapeutic options currently available for RP patients are very limited, other than one FDA approved gene therapy targeting mutations of a single gene of this highly heterogenous disease (1, 2). However, several pathogenic mechanisms, such as unfolded protein response, endoplasmic reticulum stress, mitochondrial dysfunction, oxidative stress, and abnormal disc morphogenesis, are shared between RP diseases caused by more than 3000 point mutations on ∼125 genes (RetNet, https://RetNet.org/) (3, 22, 23). Previous research suggests that some of these common signatures could be targeted to develop gene agnostic therapy for RP (3). Such pharmacological strategies include molecular chaperons stabilizing misfolded mutant protein, reducing oxidative stress, inhibiting cell death pathways and enhancing protein homeostasis (24). In many preclinical animal models of RP, dark rearing delayed degeneration while light exposure exacerbated photoreceptor loss (9). This has also been reported for the P23H mutation of rhodopsin, a common adRP in the United States (10, 11). Based on these observations we attempted to target light signaling by reducing or ablating transducin in rod or cone photoreceptors in P23H rhodopsin knock-in mice, a well-studied preclinical model of RP. Transducin, the effector G protein associated with rhodopsin or cone opsins, activates PDE6 and propagates phototransduction in the photoreceptor outer segment (13). Ablating transducin primarily blocks activation of PDE6 even in light, and mimics dark rearing in our experimental model. Our results demonstrate that reducing or ablating rod transducin delayed retinal degeneration and extended cone function in P23H mice (Figure 1, 3, 4). Additionally, reducing cone transducin preserved cone function in P23H females (Figure 5).

Dark rearing or reduction of transducin in photoreceptors is expected to elevate cGMP, metabolic rate, and oxidative stress in the retina (25–28). Still, accumulating evidence from RP models suggests that reducing rather than increasing light signaling can help to prevent blindness (9). Previous studies in a preclinical animal model of Leber’s Congenital Amaurosis (LCA) 2 demonstrated that ablating rod transducin delayed retinal degeneration (16). In this mouse model, RPE65, the enzyme involved in chromophore regeneration is ablated and the retina undergoes slow degeneration. In these mice, ablating *Gnat1* promoted photoreceptor survival by blocking constitutively active apo-opsin from initiating phototransduction cascade independent of light (16). In another RP model Rd10 mice, ablating *Gnat1* did not delay retinal degeneration (17). However, dark rearing was found to be protective in these mice, possibly through a transducin independent pathway (17). Nevertheless, our data adds to the evidence supporting targeting transducin for delaying rod degeneration and preserving cone function in a subset of inherited retinal degenerations. The potential mechanisms are discussed below.

### Protective effects of transducin; possible mechanisms

In RP patients with P23H rhodopsin mutation, rod dark adaptation is delayed more than two-fold compared to healthy adults suggesting inefficient regeneration of the chromophore bound P23H rhodopsin (29). Recent studies suggest that P23H rhodopsin can escape the ER and get transported to the rod disc membrane and has the propensity to form a homodimer (4, 30). Though such dimerization may stabilize the tertiary structure of the P23H monomer, they have reduced affinity for the chromophore 11-cis-retinal. Since apo-opsin is known to be thermally unstable (31, 32), P23H opsin without the bound chromophore could potentially be a constitutively active rhodopsin. In addition to the pathogenic mechanisms of unfolded protein response, protein aggregation and ER stress, it is possible that constitutive activation of P23H apo opsin could also contribute to photoreceptor degeneration in P23H rhodopsin RP (2, 3, 33). Previous reports of faster phototransduction activation kinetics and reduced sensitivity in P23H rods, the signatures of a constitutively active opsin, supports the possibility of this mechanism (20). Our observation of improved photoreceptor survival in P23H mice when *Gnat1* was ablated supports this hypothesis (Figure 1). Moreover, unfolded protein response and protein aggregation effect was less evident in this P23H knock-in mouse model in which abnormal disc morphogenesis was observed to be a pathogenic mechanism (6, 7). This could also explain the protective effect of dark rearing by possibly increasing the chromophore availability for the mutant protein and thus stabilizing the mutant opsin dimer. Also, rescue by dark rearing required an intact chromophore binding site within the mutant rhodopsin (11). Similar protective effect of dark rearing and *Gnat1* ablation was observed in the mouse model for LCA 2 by mitigating constitutive activation of apo-opsin (16, 34). Thus, reducing or ablating transducin can target constitutive activation of phototransduction cascade by P23H rhodopsin and promote rod survival in the degenerating retina.

Photoreceptors have a remarkably high metabolic rate and energy demand (35). Rods rely on oxidative phosphorylation in addition to aerobic glycolysis requiring high oxygen consumption rate (36, 37). Retinal oxygen consumption is higher in the dark adapted state than in the light (18) and the light-induced reduction in oxygen consumption requires phototransduction (38). Interestingly, in rd17 mice lacking functional rod transducin oxygen consumption rate was increased in the retina irrespective of the light condition possibly by simulating a constant dark-adapted state (18). This may lead to accumulation of reactive oxygen species (ROS) in rods as reported in *Gnat1^-/-^* rods (28). Such increase in oxidative stress is probably causing the reduced photoreceptor survival in our WT mice when *Gnat1* is ablated or reduced (Figure 1, Figure 2). However, in P23H retina similar oxidative stress does not seem to be detrimental to rods since we observed improved photoreceptor survival when *Gnat1* is ablated or reduced in them (Figure 1, Figure 2). During RP progression the degeneration of rods results in drastic reduction of outer retinal oxygen consumption (39, 40). This results in a significantly higher outer retinal oxygen tension as reported in P23H rat model (41), despite reduced oxygen delivery and attenuation of retinal vasculature observed in RP patients and in preclinical RP models (39, 42, 43). Exposure to such persistent high oxygen levels may result in oxidative stress in the surviving cones. When *Gnat1* is ablated in P23H retina, the higher number of surviving rods with increased oxygen consumption in their dark adapted state, may reduce the oxygen load for the surviving cones aiding in preserving their function. Additionally, the higher number of rods maintained in the P23H *Gnat1^-/-^* retina may promote cone function by secreting trophic factors that modulate metabolism and redox status of cones (44). However, the protective effect of reducing *Gnat2* on cone function in P23H retina is observed only in females suggesting sex dependent mechanisms of rescue (Figure 5).

### Sex difference in RP progression and response to *Gnat2* reduction

Epidemiological studies have reported sex dependent difference in the prevalence of retinal diseases, nevertheless such data on RP is inconclusive (45). On the contrary, in RP animal models like rd10, female mice show faster loss of rods followed by cone degeneration (46). Moreover, in the P23H mouse model, females showed accelerated loss of visual acuity, retinal thickness and ERG response and these sex specific effects on degeneration were shown to be mediated by sex hormone estradiol (14, 15). However, the mechanism of this sex hormone dependent degeneration and its specific effect on rod or cone photoreceptors are poorly understood. Our data indicates that cone function exhibited sex dependent loss of ERG response in females whereas such sex difference was less apparent in rod function loss (Figure 4). Since our experiments on rod ERG response were done on mice lacking cone phototransduction, we speculate that cone function is exacerbating sex dependent degeneration in P23H females. Even though oxidative damage and reduced availability of glucose has been suggested to cause cone death in RP (47, 48) it is not known how these mechanisms are different in male and female retinas resulting in sex hormone dependent RP progression in females. It is important to note that male and female retinas are inherently different in terms of their transcriptomic, proteomic and metabolic signatures, besides the presence of systemic and locally secreted sex hormones (49–51).

Additionally, our data show that the improvement in cone function by reducing *Gnat2* was observed only in females, while that in rod function by reducing *Gnat1* was observed in both sexes. This rescued P23H female cone function was comparable to that of P23H males suggesting that *Gnat2* reduction compensated for the accelerated degenerating mechanism in females. Notably, cone photoreceptors do not get saturated in bright light and therefore maintain high ATP consumption even in light to facilitate light response (27). Such high energy demand coupled with higher oxygen tension and altered metabolic status in RP may lead to higher oxygen consumption and oxidative stress in P23H cones. Downregulating *Gnat2* may reduce cone phototransduction and may in turn reduce cone oxygen consumption. Moreover, cone light response amplitude was reduced in WT Gnat2^+/-^ mice (Figure S6). Thus, improvement of female cone function by *Gnat2* downregulation is possibly through modulating cellular redox status and metabolic activity of P23H cones (52). Additionally, it is possible that the sex hormone mediated degeneration of female cones is through increased oxidative stress. Interestingly, tamoxifen, a selective estrogen receptor modulator rescued cone function in a NaIO3 model of oxidative stress induced retinal degeneration (53). In another study, tamoxifen was protective against retinal degeneration in rd10 model of RP and in the light induced degeneration (54). Similarly, in a P23H rat model therapeutic effect of reserpine in delaying RP progression was only observed in females (50). In this case, reserpine treatment altered expression levels of genes involved in proteostasis and cellular stress. The pathogenic mechanisms and the effectiveness of therapies may vary between sexes since females display accelerated photoreceptor degeneration and loss of visual function in many RP models including P23H rhodopsin mutants (14, 15, 46). More experiments are needed to elucidate how inherent differences in male and female retina interact with RP progression pathways and homeostatic processes promoting cell survival, in rods and cones, to inform future therapeutic approaches. Nevertheless, our data points to potential sex specific therapies for delaying RP progression.

Overall, the data presented here indicates that transducin downregulation is a potential therapeutic strategy for delaying rod degeneration and preserving cone function in P23H rhodopsin RP. Based on previous reports this strategy may be applied to a broad subset of RP for delaying photoreceptor degeneration. Additionally, our data provides better insights into the mechanisms of photoreceptor degeneration in RP, paving the way for future studies to identify effective therapies to preserve vision in RP patients.

## METHODS

### Preclinical models

All animal experimental protocols adhered to Guide for the Care and Use of Laboratory Animals and were approved by the Institutional Animal Care and Use Committee at the University of Utah. Mice carrying the heterozygous P23H mutation in rhodopsin (P23H mice) were generated by crossing *Rho^P23H/P23H^* males (6) with C57Bl/6J female mice (The Jackson Laboratory, stock # 000664). Experimental P23H mice and littermate WT mice were generated using *Rho^P23H/WT^* and *Rho^WT/WT^* breeding pairs. We generated P23H mice lines on *Gnat1^-/-^* (19) and *Gnat2^-/-^* (55) background by crossing P23H mice with *Gnat1^-/-^* (The Jackson Laboratory, stock #) or *Gnat2^-/-^* mice, a kind gift from Dr. Marie Burns, University of California, Davis. The experimental mice lacking functional cones, P*23H Gnat2^-/-^*, *P23H Gnat2^-/-^ Gnat1^+/-^*, *WT Gnat2^-/-^* and *WT Gnat2^-/-^ Gnat1^+/-^*were generated using P*23H Gnat2^-/-^* and *WT Gnat2^-/-^ Gnat1*^+/-^ breeding pair. The experimental mice lacking functional rods P*23H Gnat1^-/-^*, P*23H Gnat1^-/-^ Gnat2^+/-^*, *WT Gnat1^-/-^* and *WT Gnat1^-/-^ Gnat2^+/-^*were generated using P*23H Gnat1^-/-^* and *WT Gnat1^-/-^ Gnat2*^+/-^ breeding pair. Both male and female mice were used in this study. Mice were kept under 12/12 hour light/dark cycle with 300 lux light intensity during the light phase with free access to food and water.

### Optical Coherence Tomography

Mice were anesthetized with ketamine (100 mg/kg) (MWI Animal Health) and xylazine (10 mg/kg) (VET One) by intraperitoneal injections, and their pupils were dilated with 1% Atropine Sulfate (Amneal Pharmaceuticals). Eyes were kept moist using Hypromellose (2.5%) ophthalmic solution (Vista Gonio). OCT b-scan images were acquired using Micron IV (Phoenix) and ONL thickness was measured at 300μm from the Optic Nerve Head (ONH) from superior and inferior quadrants using manual calipers. Two B scans from each eye were used for measurement, and the average of all the calipers of each eye is used for the analysis.

### Ex vivo Electroretinogram

Transretinal (ex vivo) ERG was done as described previously (56). Briefly, mice were dark adapted for 24 hours and euthanized by CO2 and cervical dislocation under dim red light. The enucleated eyes were immediately placed into carbonated (95% O2/5% CO2) Ames’ medium (A1420; Sigma-Aldrich) containing 1.932 g/L NaHCO3 (Sigma-Aldrich). The retina was removed from the underlying RPE/choroid and mounted on the ex vivo ERG specimen holder. The mounted retina was perfused at a rate of 1 mL/min with heated Ames’ medium (34°C) containing 100µM BaCl2 (Sigma-Aldrich) to eliminate the Müller glia component of the ERG signal. From the dark-adapted retina, ERG responses to flashes of increasing intensity of light (530 nm, 2–10 ms flash duration) were recorded. To isolate the photoreceptor component of the ERG signal, 40 µM DL-AP4 (Cat. #0101; Tocris Bioscience) was added to the perfusion medium. ON bipolar cell responses were determined by subtracting the photoreceptor response from the ERG response. To determine flash intensity (I) required to generate half-maximal photoreceptor (I_1/2_), a Hill function

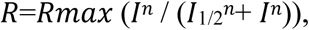

was fitted to amplitude (R) data using the GraphPad Prism nonlinear curve fit tool. Rmax is the maximal response amplitude and n is Hill’s coefficient.

### Immunohistochemistry

Eyes were enucleated and immediately fixed in ice-cold 4% paraformaldehyde (PFA) for 20 minutes on ice. After rinsing with 1 X phosphate buffered saline (PBS) lenses were removed and the eyecups were cryoprotected in a sucrose gradient (10 %, 20 %, 30 %). These were placed in a 1:2 solution of 20 % sucrose/PBS and OCT (Sakura Finetek) mix overnight at 4°C and embedded in 20 % sucrose/PBS and OCT mix and cryo-sections were made at 10μm thickness. Slides were rinsed 3 x 10 minutes in PBS to remove OCT. Non-specific binding was blocked for one hour at room temperature with blocking buffer made of 0.1 % Triton-X 100 (Fisher Scientific) and 10 % normal goat serum in PBS. Samples were incubated with 1:100 dilution of Cy5 conjugated PNA (Vector Laboratories, CL-1075-1) for 1 hour in a humidified dark chamber at 4 °C. The slides were rinsed for 3 x 10 minutes in PBS, then incubated with Hoescht 33342 (Invitrogen, Cat. No: H3570) for 30 minutes at room temperature. Coverslips were mounted with Fluoromount-G (SouthernBiotech, Cat. No.: 0100-10) mounting media. Images of retinal sections were obtained using a Zeiss LSM800 confocal microscope and the number of cones in 100μm blocks in the central retina (500μm from ONH) were quantified using ImageJ software (v1.54p).

### Statistics

Statistical analysis was performed by GraphPad Prism (v10). Analysis was done by 2-way ANOVA, followed by Tukey’s multiple-comparison test. ERG responses to multiple flashes were analyzed by 2-way ANOVA between groups, with the genotype/sex as factor 1 and flash intensity as factor 2. The groups were identified to be different from each other when *P* < 0.05. Data are shown as mean ± SEM.

## Acknowledgements

This work was supported by NIH R01 (EY034986) to FV and P30 (EY014800), T32 (EY024234) and Unrestricted Grant from Research to Prevent Blindness to Moran Eye Center, University of Utah. Authors thank Prof. Marie Burns, University of California, Davis for kindly providing Gnat2^-/-^ mice.

## Author contributions

DM and FV conceptualized the study and developed methodology. DM, LS, and MM performed the investigation. DM wrote the original draft of the manuscript. DM and FV reviewed and edited the manuscript.

---

**Figure S1.**
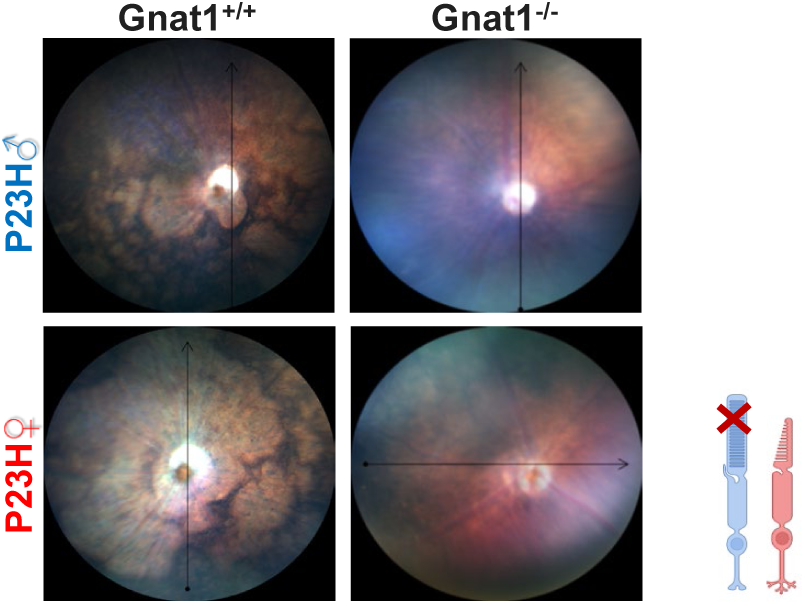
Abolishing rod transducin is protective in P23H mice. Fundus images from 12-month-old *P23H Gnat1^-/-^* male and female mice did not show the characteristic appearance of advanced Retinitis Pigmentosa as exhibited in *P23H Gnat1^+/+^*.

**Figure S2.**
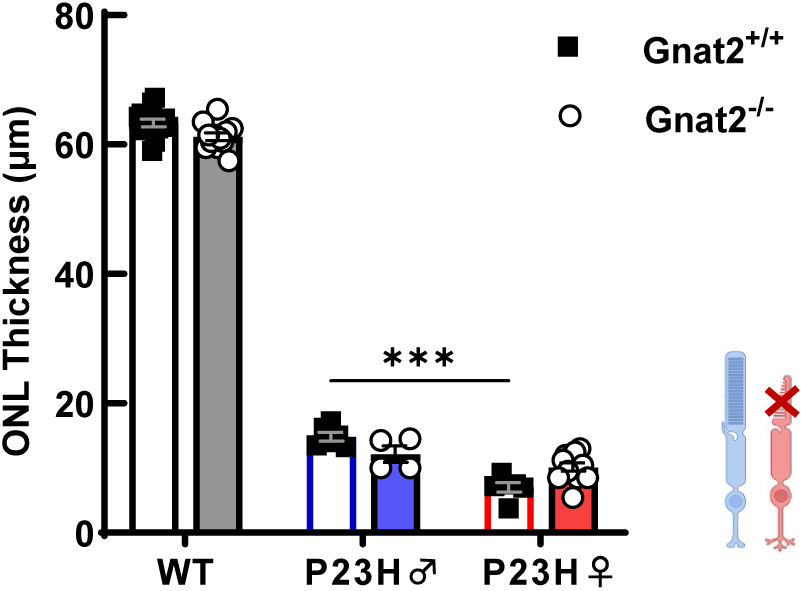
Ablating cone transducin did not affect photoreceptor survival in P23H mice: Quantification of ONL thickness of 7-month-old P23H males and females compared with those with their *Gnat2* ablated, showing unaltered photoreceptor number. Tukey’s post hoc tests, ***p<0.001. Mean ± SEM, N=4-10

**Figure S3.**
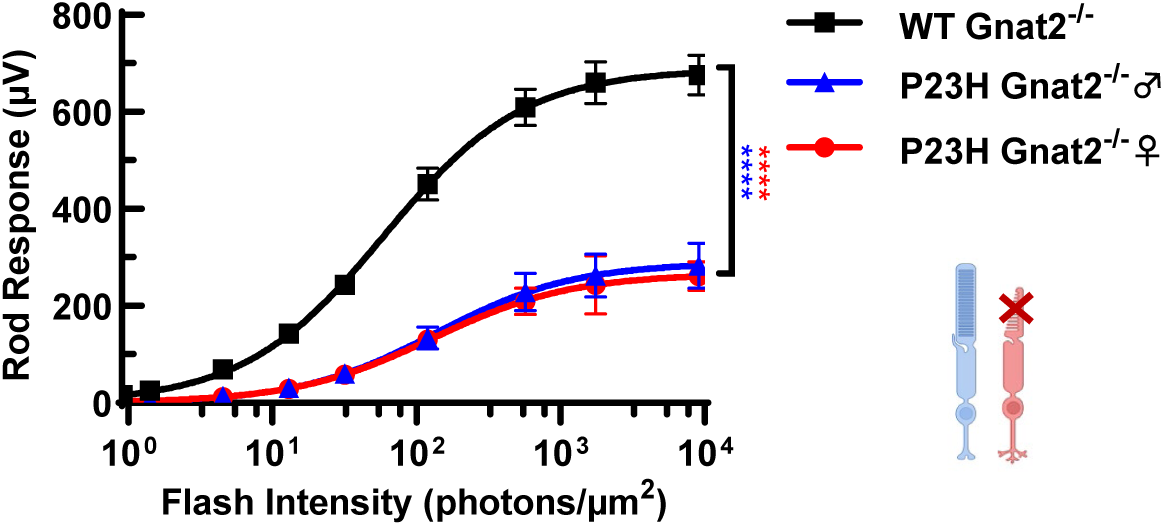
Rod light response were equally reduced in P23H males and females. Ex vivo ERG amplitude data as a function of light flash intensity from 3-month-old *WT Gnat2^-^*^/-^ and *P23H Gnat2^-/-^* males and females showing equally reduced rod light response in both sexes. Smooth traces are Hill’s equation fitted to the mean amplitude data. Mean ± SEM; two-way ANOVA, ****p<0.0001. N=4-7

**Figure S4.**
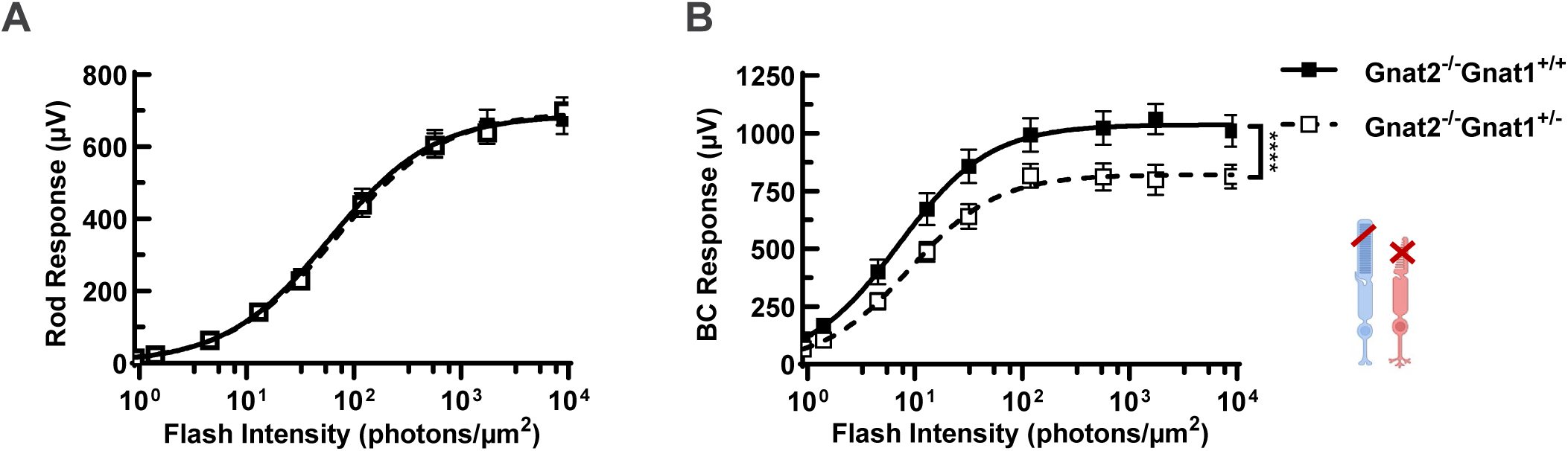
Downregulating rod transducin reduced rod mediated bipolar cell response in WT mice. Ex vivo ERG amplitude data as a function of light flash intensity from 3-month-old *WT Gnat2^-/-^ Gnat1^+/+^* and *WT Gnat2^-/-^ Gnat1^+/-^* ; rod response (A), rod mediated bipolar response (B), demonstrating suppressed rod to rod bipolar cell signaling when *Gnat1* expression was reduced. Mean ± SEM; two-way ANOVA, *p<0.05, **p<0.01, ***p<0.001. Smooth traces in (A-B) are Hill’s equation fitted to the mean amplitude data. Mean ± SEM, N=4-7

**Figure S5.**
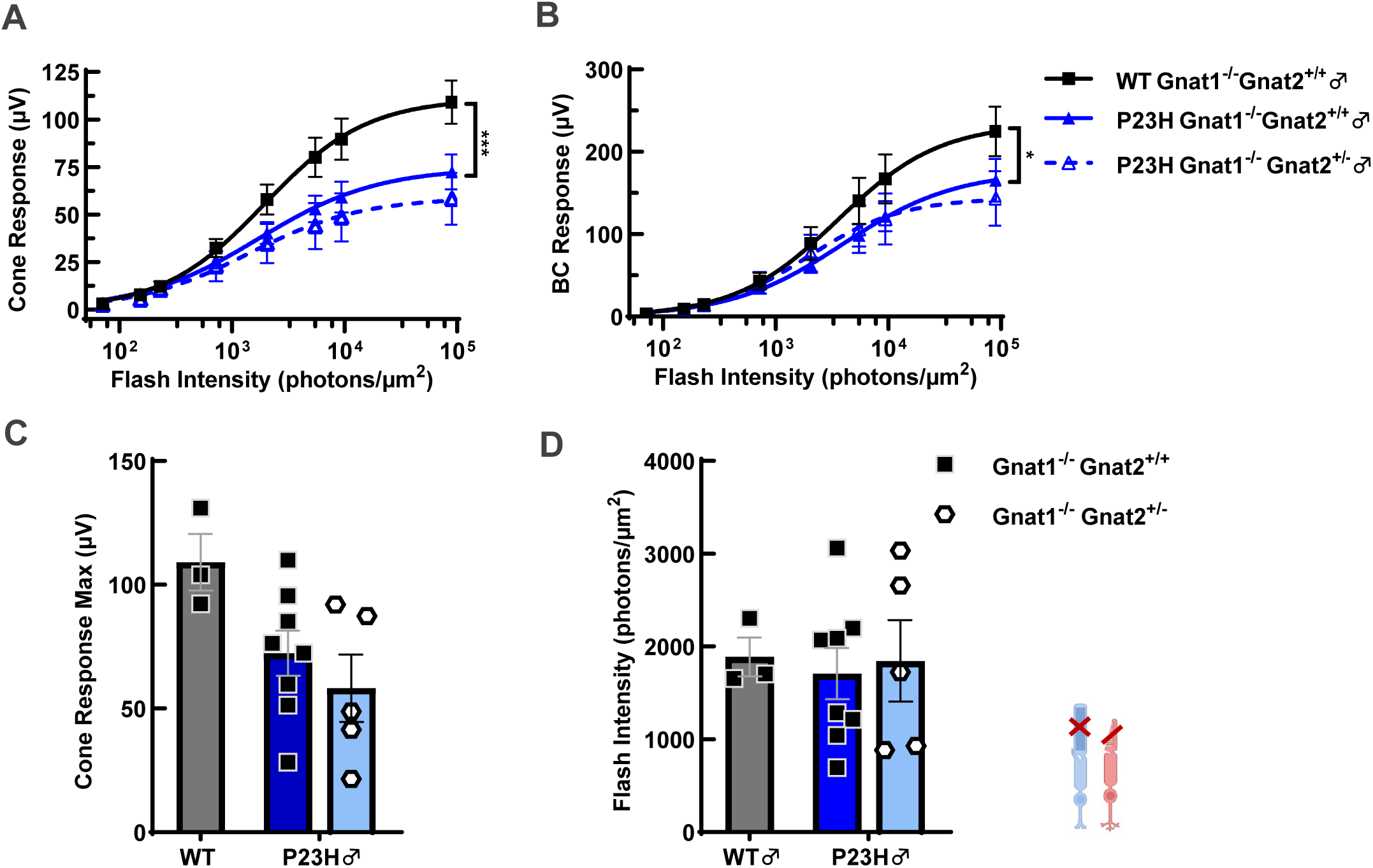
Reducing cone transducin did not improve cone function in P23H males: Cone ERG Response to increasing intensities of light flashes from 12-14 months old WT *Gnat1^-/-^* and *P23H Gnat1^-/-^* males compared with *P23H Gnat1^-/-^* males carrying a single allele of *Gnat2* demonstrating unaltered cone photoreceptor response (A) and bipolar cell response (B). Smooth traces in (A and B) are Hill’s equation fitted to the mean amplitude data. Two-way ANOVA, *p<0.05, ***p<0.001. No significant difference was observed in maximum cone response amplitude (C) or cone light sensitivity determined by the intensity required to generate half-maximal cone response (D). Mean ± SEM, N=3-8

**Figure S6.**
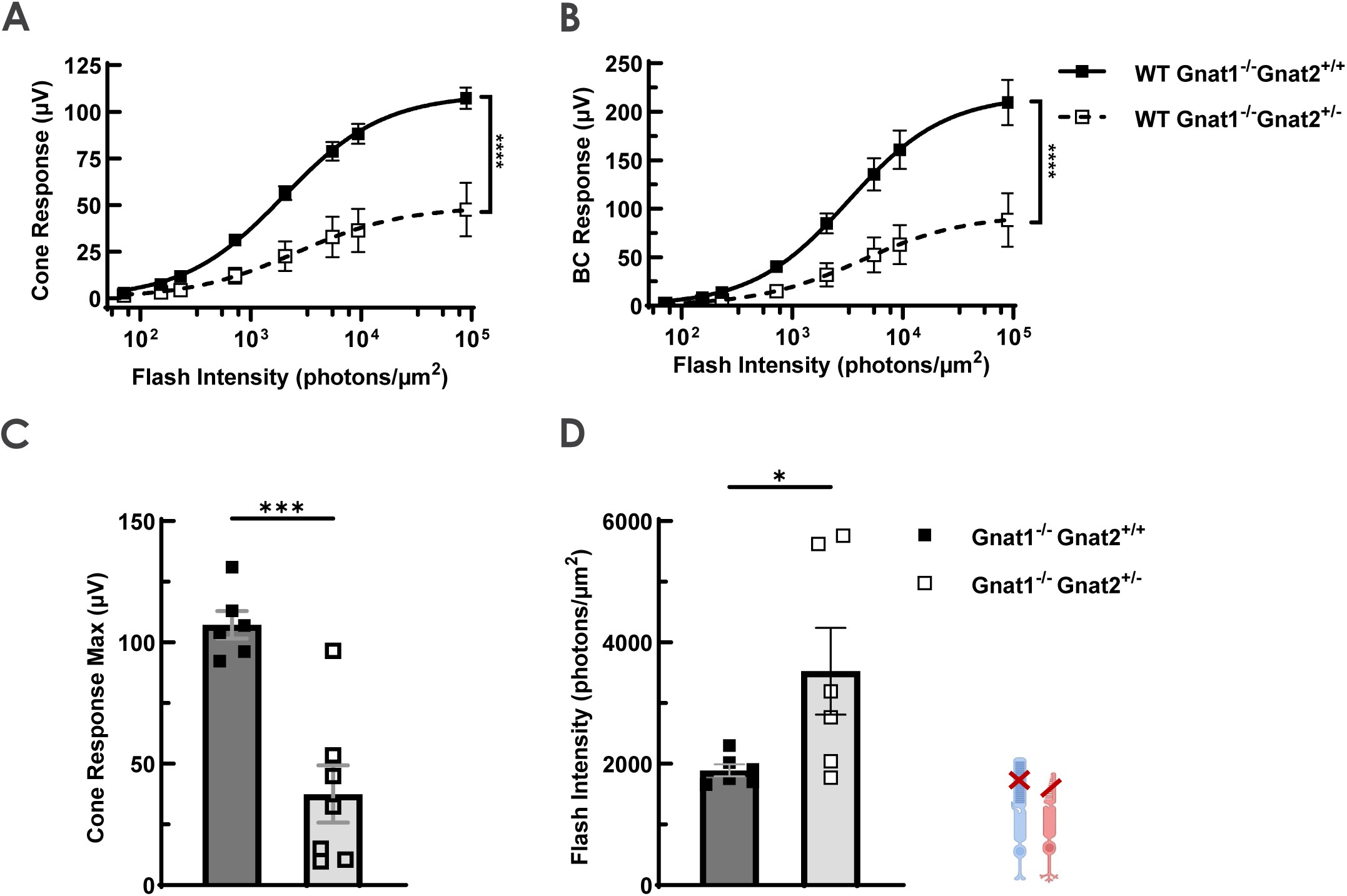
Reducing cone transducin reduced cone function in older WT mice: Cone ERG Response from 12-14 months old *P23H Gnat1^-/-^* mice compared with those carrying a single allele of *Gnat2* showing drastically reduced cone photoreceptor response (A) and bipolar cell response (B). Smooth traces in (A and B) are Hill’s equation fitted to the mean amplitude data. Two-way ANOVA, ****p<0.0001. Maximum cone response amplitude (C) and cone light sensitivity determined by the intensity required to generate half-maximal cone response (D) was also reduced when *Gnat2* was heterozygous in *P23H Gnat1^-/-^* mice. Tukey’s post hoc tests, *p<0.05, ***p<0.001. Mean ± SEM, N=6-7.

**Figure S7.**
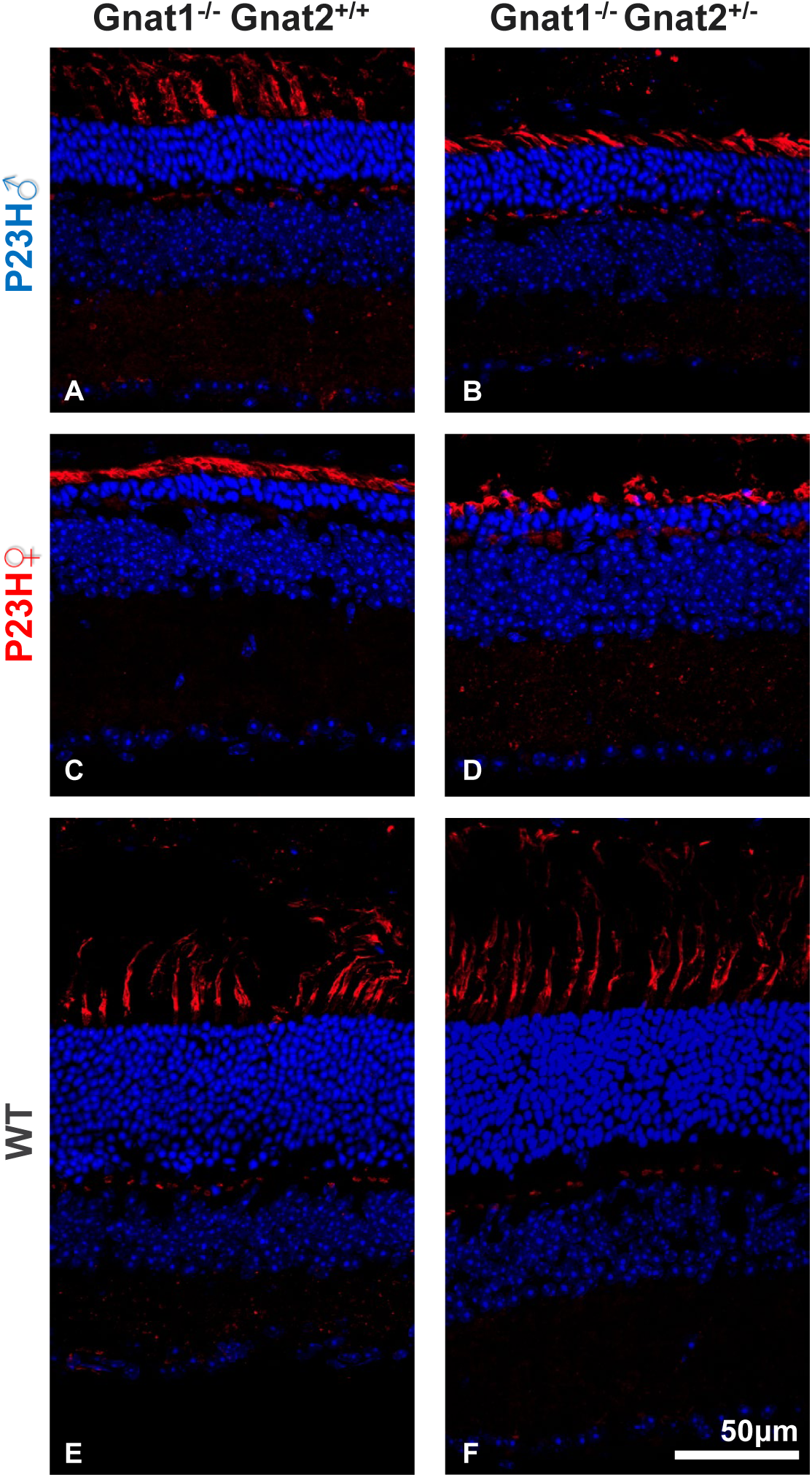
Reducing *Gnat2* did not affect cone photoreceptor survival in 12–14-month P23H mice. Representative confocal images of retina sections stained with PNA (red) for cones and Hoechst (blue) for nuclei from *P23H Gnat1^-/-^ Gnat2^+/+^* males (A) *P23H Gnat1^-/-^ Gnat2^+/-^* males (B) *P23H Gnat1^-/-^ Gnat2^+/+^* females (C), *P23H Gnat1^-/-^ Gnat2^+/-^* females (D), WT *Gnat1^-/-^ Gnat2^+/+^* (E) and WT *Gnat1^-/-^ Gnat2^+/-^* mice (F).

## REFERENCES

1. Schneider N, Sundaresan Y, Gopalakrishnan P, Beryozkin A, Hanany M, Levanon EY, et al. Inherited retinal diseases: Linking genes, disease-causing variants, and relevant therapeutic modalities. Progress in Retinal and Eye Research. 2022;89:101029.

2. Tolone A, Şen M, Chen Y, Ueffing M, Arango-Gonzalez B, and Paquet-Durand F. Pathomechanisms of Inherited Retinal Degeneration and Perspectives for Neuroprotection. Cold Spring Harbor Perspectives in Medicine. 2022;13:a041310.

3. Athanasiou D, Aguila M, Bellingham J, Li W, McCulley C, Reeves PJ, and Cheetham ME. The molecular and cellular basis of rhodopsin retinitis pigmentosa reveals potential strategies for therapy. Prog Retin Eye Res. 2018;62:1–23.

4. Woods KN, and Pfeffer J. Conformational perturbation, allosteric modulation of cellular signaling pathways, and disease in P23H rhodopsin. Scientific Reports. 2020;10(1):2657.

5. Miller LM, Gragg M, Kim TG, and Park PS-H. Misfolded opsin mutants display elevatedβ-sheet structure. FEBS Letters. 2015;589(20PartB):3119–25.

6. Sakami S, Maeda T, Bereta G, Okano K, Golczak M, Sumaroka A, et al. Probing Mechanisms of Photoreceptor Degeneration in a New Mouse Model of the Common Form of Autosomal Dominant Retinitis Pigmentosa due to P23H Opsin Mutations. Journal of Biological Chemistry. 2011;286(12):10551–67.

7. Sakami S, Kolesnikov AV, Kefalov VJ, and Palczewski K. P23H opsin knock-in mice reveal a novel step in retinal rod disc morphogenesis. Human Molecular Genetics. 2013;23(7):1723–41.

8. Chiang WC, Kroeger H, Sakami S, Messah C, Yasumura D, Matthes MT, et al. Robust Endoplasmic Reticulum-Associated Degradation of Rhodopsin Precedes Retinal Degeneration. Mol Neurobiol. 2015;52(1):679–95.

9. Paskowitz DM, LaVail MM, and Duncan JL. Light and inherited retinal degeneration. Br J Ophthalmol. 2006;90(8):1060–6.

10. Naash ML, Peachey NS, Li ZY, Gryczan CC, Goto Y, Blanks J, et al. Light-induced acceleration of photoreceptor degeneration in transgenic mice expressing mutant rhodopsin. Invest Ophthalmol Vis Sci. 1996;37(5):775–82.

11. Tam BM, and Moritz OL. Dark Rearing Rescues P23H Rhodopsin-Induced Retinal Degeneration in a Transgenic Xenopus laevis Model of Retinitis Pigmentosa: A Chromophore-Dependent Mechanism Characterized by Production of N-Terminally Truncated Mutant Rhodopsin. The Journal of Neuroscience. 2007;27(34):9043–53.

12. Brill E, Malanson KM, Radu RA, Boukharov NV, Wang Z, Chung HY, et al. A novel form of transducin-dependent retinal degeneration: accelerated retinal degeneration in the absence of rod transducin. Invest Ophthalmol Vis Sci. 2007;48(12):5445–53.

13. Arshavsky VY, and Burns ME. Photoreceptor Signaling: Supporting Vision across a Wide Range of Light Intensities. Journal of Biological Chemistry. 2012;287(3):1620–6.

14. Barwick SR, Xiao H, Wolff D, Wang J, Perry E, Marshall B, and Smith SB. Sigma 1 receptor activation improves retinal structure and function in the RhoP23H/+ mouse model of autosomal dominant retinitis pigmentosa. Experimental Eye Research. 2023;230:109462.

15. Rowe AA, Velasquez MJ, Aumeier JW, Reyes S, Yee T, Nettesheim ER, et al. Female sex hormones exacerbate retinal neurodegeneration. Science Advances. 2025;11(15):eadr6211.

16. Woodruff ML, Wang Z, Chung HY, Redmond TM, Fain GL, and Lem J. Spontaneous activity of opsin apoprotein is a cause of Leber congenital amaurosis. Nature Genetics. 2003;35(2):158–64.

17. Sundar JC, Munezero D, Bryan-Haring C, Saravanan T, Jacques A, and Ramamurthy V. Rhodopsin signaling mediates light-induced photoreceptor cell death in rd10 mice through a transducin-independent mechanism. Human Molecular Genetics. 2019;29(3):394–406.

18. Berkowitz BA, Podolsky RH, Childers KL, Roberts R, Katz R, Waseem R, et al. Transducin-Deficient Rod Photoreceptors Evaluated With Optical Coherence Tomography and Oxygen Consumption Rate Energy Biomarkers. Investigative Ophthalmology & Visual Science. 2022;63(13):22-.

19. Calvert PD, Krasnoperova NV, Lyubarsky AL, Isayama T, Nicoló M, Kosaras B, et al. Phototransduction in transgenic mice after targeted deletion of the rod transducin α-subunit. Proceedings of the National Academy of Sciences. 2000;97(25):13913–8.

20. Pasquale RL, Guo Y, Umino Y, Knox B, and Solessio E. Temporal Contrast Sensitivity Increases despite Photoreceptor Degeneration in a Mouse Model of Retinitis Pigmentosa. eneuro. 2021;8(2):ENEURO.0020–21.2021.

21. Majumder A, Pahlberg J, Boyd KK, Kerov V, Kolandaivelu S, Ramamurthy V, et al. Transducin translocation contributes to rod survival and enhances synaptic transmission from rods to rod bipolar cells. Proc Natl Acad Sci U S A. 2013;110(30):12468–73.

22. Zhen F, Zou T, Wang T, Zhou Y, Dong S, and Zhang H. Rhodopsin-associated retinal dystrophy: Disease mechanisms and therapeutic strategies. Front Neurosci. 2023;17:1132179.

23. SP Daiger BR, J Greenberg, A Christoffels, W Hide. Investigative Ophthalmology & Visual Science. 1998:S295.

24. Chew LA, and Iannaccone A. Gene-agnostic approaches to treating inherited retinal degenerations. Front Cell Dev Biol. 2023;11:1177838.

25. Tolone A, Belhadj S, Rentsch A, Schwede F, and Paquet-Durand F. The cGMP Pathway and Inherited Photoreceptor Degeneration: Targets, Compounds, and Biomarkers. Genes (Basel). 2019;10(6).

26. Okawa H, Sampath A, Laughlin S, and Fain G. ATP Consumption by Mammalian Rod Photoreceptors in Darkness and in Light. Current biology : CB. 2008;18:1917–21.

27. Ingram NT, Fain GL, and Sampath AP. Elevated energy requirement of cone photoreceptors. Proceedings of the National Academy of Sciences. 2020;117(32):19599–603.

28. Liu H, Tang J, Du Y, Saadane A, Samuels I, Veenstra A, et al. Transducin1, Phototransduction and the Development of Early Diabetic Retinopathy. Investigative Ophthalmology & Visual Science. 2019;60(5):1538–46.

29. Kemp CM, Jacobson SG, Roman AJ, Sung C-H, and Nathans J. Abnormal Rod Dark Adaptation in Autosomal Dominant Retinitis Pigmentosa With Proline-23-Histidine Rhodopsin Mutation. American Journal of Ophthalmology. 1992;113(2):165–74.

30. Chen Y, Jastrzebska B, Cao P, Zhang J, Wang B, Sun W, et al. Inherent Instability of the Retinitis Pigmentosa P23H Mutant Opsin. Journal of Biological Chemistry. 2014;289(13):9288–303.

31. Cornwall MC, and Fain GL. Bleached pigment activates transduction in isolated rods of the salamander retina. The Journal of Physiology. 1994;480(2):261–79.

32. Zhukovsky EA, Robinson PR, and Oprian DD. Transducin Activation by Rhodopsin Without a Covalent Bond to the 11-Cis-Retinal Chromophore. Science. 1991;251(4993):558–60.

33. Fain GL. Why photoreceptors die (and why they don’t). BioEssays. 2006;28(4):344–54.

34. Fan J, Woodruff ML, Cilluffo MC, Crouch RK, and Fain GL. Opsin activation of transduction in the rods of dark-reared Rpe65 knockout mice. J Physiol. 2005;568(Pt 1):83–95.

35. Okawa H, Sampath AP, Laughlin SB, and Fain GL. ATP Consumption by Mammalian Rod Photoreceptors in Darkness and in Light. Current Biology. 2008;18(24):1917–21.

36. Kanow MA, Giarmarco MM, Jankowski CSR, Tsantilas K, Engel AL, Du J, et al. Biochemical adaptations of the retina and retinal pigment epithelium support a metabolic ecosystem in the vertebrate eye. eLife. 2017;6:e28899.

37. Chen Y, Zizmare L, Calbiague V, Wang L, Yu S, Herberg FW, et al.: eLife Sciences Publications, Ltd; 2024.

38. Du J, Rountree A, Cleghorn WM, Contreras L, Lindsay KJ, Sadilek M, et al. Phototransduction Influences Metabolic Flux and Nucleotide Metabolism in Mouse Retina. J Biol Chem. 2016;291(9):4698–710.

39. Rahimi M, Leahy S, Matei N, Blair NP, Jeong S, Craft CM, and Shahidi M. Assessment of inner retinal oxygen metrics and thickness in a mouse model of inherited retinal degeneration. Experimental Eye Research. 2021;205:108480.

40. Türksever C, Valmaggia C, Orgül S, Schorderet DF, Flammer J, and Todorova MG. Retinal vessel oxygen saturation and its correlation with structural changes in retinitis pigmentosa. Acta Ophthalmologica. 2014;92(5):454–60.

41. Yu D-Y, and Cringle SJ. Retinal degeneration and local oxygen metabolism. Experimental Eye Research. 2005;80(6):745–51.

42. Ma Y, Kawasaki R, Dobson LP, Ruddle JB, Kearns LS, Wong TY, and Mackey DA. Quantitative Analysis of Retinal Vessel Attenuation in Eyes with Retinitis Pigmentosa. Investigative Ophthalmology & Visual Science. 2012;53(7):4306–14.

43. Fernández-Sánchez L, Esquiva G, Pinilla I, Lax P, and Cuenca N. Retinal Vascular Degeneration in the Transgenic P23H Rat Model of Retinitis Pigmentosa. Front Neuroanat. 2018;12:55.

44. Clérin E, Aït-Ali N, Sahel JA, and Léveillard T. Restoration of Rod-Derived Metabolic and Redox Signaling to Prevent Blindness. Cold Spring Harb Perspect Med. 2024;14(11).

45. Tillmann A, Ceklic L, Dysli C, and Munk MR. Gender differences in retinal diseases: A review. Clinical and Experimental Ophthalmology. 2024;52(3):317–33.

46. Li B, Gografe S, Munchow A, Lopez-Toledano M, Pan Z-H, and Shen W. Sex-related differences in the progressive retinal degeneration of the rd10 mouse. Experimental Eye Research. 2019;187:107773.

47. Campochiaro PA, and Mir TA. The mechanism of cone cell death in Retinitis Pigmentosa. Progress in Retinal and Eye Research. 2018;62:24–37.

48. Lee SJ, Emery D, Vukmanic E, Wang Y, Lu X, Wang W, et al. Metabolic transcriptomics dictate responses of cone photoreceptors to retinitis pigmentosa. Cell Reports. 2023;42(9).

49. Jang GF, Crabb JS, Grenell A, Wolk A, Campla C, Luo S, et al. Quantitative proteomic profiling reveals sexual dimorphism in the retina and RPE of C57BL6 mice. Biol Sex Differ. 2024;15(1):87.

50. Song HB, Campello L, Mondal AK, Chen HY, English MA, Glen M, et al. Sex-specific attenuation of photoreceptor degeneration by reserpine in a rhodopsin P23H rat model of autosomal dominant retinitis pigmentosa. 2025.

51. Saravanan M, Xu R, Roby O, Wang Y, Zhu S, Lu A, and Du J. Tissue-Specific Sex Difference in Mouse Eye and Brain Metabolome Under Fed and Fasted States. Invest Ophthalmol Vis Sci. 2023;64(3):18.

52. Punzo C, Xiong W, and Cepko CL. Loss of Daylight Vision in Retinal Degeneration: Are Oxidative Stress and Metabolic Dysregulation to Blame?. Journal of Biological Chemistry. 2012;287(3):1642–8.

53. Lee TT, Bell BA, Anderson BD, Song Y, and Dunaief JL. Tamoxifen protects photoreceptors in the sodium iodate model. Experimental Eye Research. 2024;242:109879.

54. Wang X, Zhao L, Zhang Y, Ma W, Gonzalez SR, Fan J, et al. Tamoxifen Provides Structural and Functional Rescue in Murine Models of Photoreceptor Degeneration. The Journal of Neuroscience. 2017;37(12):3294–310.

55. Ronning K, Peinado Allina G, Miller E, Zawadzki R, Pugh JE, Hermann R, and Burns M. Loss of cone function without degeneration in a novel Gnat2 knock-out mouse. Experimental Eye Research. 2018;171.

56. Vinberg F, and Kefalov V. Simultaneous ex vivo functional testing of two retinas by in vivo electroretinogram system. J Vis Exp. 2015(99):e52855.

